# Preserved brain function and reversible cognitive adaptation during endurance exercise

**DOI:** 10.64898/2026.04.16.719122

**Authors:** Iñigo López, Soraya Pozueta, Izaskun Yurrebaso-Santamaría, Eneritz López-Muguruza, Gixane González-García, Carlos Matute

## Abstract

Endurance exercise imposes extreme metabolic demands on the adult human brain, raising the question of how core brain function is preserved under physiological challenge. We previously showed that marathon running induces reversible reductions in myelin within specific white-matter tracts, suggesting adaptive structural change under metabolic stress. Here, we asked whether this process is functionally tolerated. Neurophysiological recordings revealed maintained conduction latencies across motor, somatosensory, visual, and auditory pathways within 48 hours after race completion, indicating intact axonal signal transmission despite reduced myelin content. Cognitive testing revealed selective and transient modulation of higher-order processing, including attenuated practice-related gains in processing speed and short-lived increases in interference, whereas visuomotor speed and executive flexibility were preserved. All cognitive measures normalized one month after the race, supporting an adaptive framework linking myelin change with preserved brain function under extreme metabolic stress.

## Main Text

Prolonged endurance exercise imposes extreme metabolic demands that require coordinated adaptations across multiple organs to sustain performance^1^. During marathon running, carbohydrates constitute the primary energy source, but as glycogen stores become progressively depleted in muscle, liver, and other tissues, metabolism shifts toward lipid utilization. Fat provides a more abundant and energetically dense substrate than carbohydrates, enabling sustained energy supply during prolonged exertion^2^. Although the metabolic consequences of endurance exercise are well characterized in peripheral organs, how the human brain adapts to sustained energetic stress remains poorly understood.

The brain operates under high and continuous energy demand and relies predominantly on glucose under resting conditions. However, accumulating evidence indicates that under conditions of increased demand or limited glucose availability, alternative substrates can contribute to neural energy metabolism^3^. In particular, fatty acids have recently emerged as important modulators of brain function, influencing not only membrane structure but also electrophysiological processes, synaptic transmission, and cognition through effects on ion-channel function and receptor-mediated signaling^4^. These observations challenge the traditional view that neuronal metabolism is exclusively glucose dependent and suggest a broader metabolic flexibility in the adult brain.

Myelin, which enwraps axons in both the central and peripheral nervous systems, is exceptionally lipid rich, with lipids accounting for 70–80% of its mass^5^. Beyond its classical role in electrical insulation and saltatory conduction, myelin provides metabolic support to axons, and oligodendroglial lipid metabolism has been proposed to contribute to neural energy homeostasis^6^. Together with emerging evidence for functional roles of fatty acids in neural physiology, these findings raise the possibility that myelin represents a strategically positioned lipid reservoir within the brain, analogous to peripheral fat stores mobilized during prolonged exercise. In support of this idea, we recently demonstrated that marathon running induces a reversible reduction of myelin content in specific white-matter tracts engaged during locomotion, sensorimotor integration, and emotional processing^7^. The spatial selectivity and reversibility of these changes suggested an adaptive physiological process rather than structural damage. Whether such exercise-induced myelin remodeling is functionally tolerated by the adult human brain, preserving neural signal transmission and cognition under extreme metabolic stress, remains unknown.

Here, we directly addressed this question by combining neurophysiological and cognitive assessments in human marathon runners. We tested whether transient reductions in myelin content are associated with alterations in axonal signal transmission across motor and sensory pathways using evoked potential measurements spanning central and peripheral circuits. We further examined whether endurance exercise is accompanied by changes in cognitive performance, focusing on tasks probing processing speed, cognitive interference, and executive flexibility. By integrating functional readouts with prior structural findings, this study provides a proof-of-principle examination of how metabolic adaptations in the adult human brain relate to neural signaling and cognitive performance under extreme physiological challenge.

### Myelin reduction in endurance exercise conserves axonal signal transmission

Having established by MRI that marathon running induces a reversible reduction of myelin content in white-matter tracts engaged during motor, sensory, and integrative processing^7^, we next asked whether this adaptive structural remodeling is functionally tolerated by the adult human brain. Because myelin is classically regarded as a principal determinant of axonal conduction velocity, even transient changes in myelin content would be predicted to alter the timing of neural signal propagation^8^. To test this prediction, we measured conduction latencies across central motor and sensory pathways before and after marathon running.

Motor-evoked potentials elicited by transcranial magnetic stimulation of the motor cortex provided a functional readout of corticospinal signal transmission to upper- and lower-limb muscles^9^. Latencies recorded from median and tibial nerves were maintained across all sessions, with no differences between pre-race measurements, recordings obtained within 48 hours after race completion, or follow-up assessments one month later (Fig. 1A, B). These findings indicate that axonal signal transmission along motor pathways remains intact despite exercise-induced myelin remodeling. We next assessed sensory pathway conduction using cortical somatosensory evoked potentials, which reflect the timing and integrity of ascending afferent signals^10^. Peak latencies of the lower-limb P37 component following posterior tibial nerve stimulation and the upper-limb N20 component following median nerve stimulation were unchanged across all time points (Fig. 1C, D), consistently across left and right limbs.

**Figure 1.**
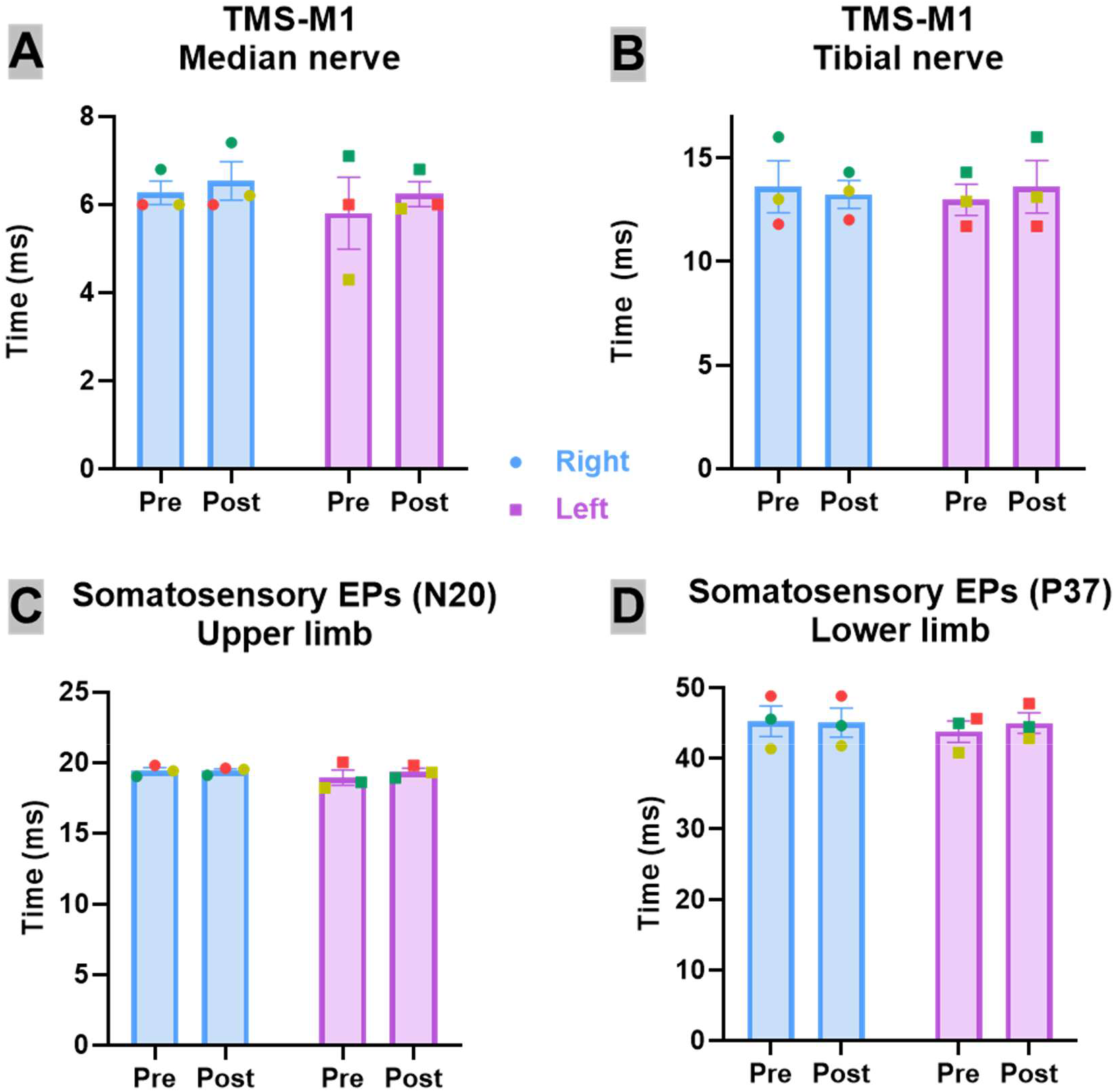
Maintained neural signal transmission after endurance exercise despite transient myelin remodeling. Neurophysiological recordings revealed stable conduction latencies across motor and somatosensory pathways within 48 h after marathon completion, indicating unchanged pathway conduction timing (no detectable change in evoked-potential latencies) despite reduced myelin content. Motor-evoked potentials elicited by transcranial magnetic stimulation showed stable latencies in median (A) and tibial (B) nerve pathways across pre- and post-race. Likewise, cortical somatosensory evoked potentials revealed unchanged N20 (upper limb) and P37 (lower limb) peak latencies (C, D), with consistent results across hemispheres and time points. Together, these data demonstrate conserved axonal signal fidelity under extreme metabolic demand. n = 3, no consistent within-subject shifts, paired two-tailed t test, in all comparisons. Color codes represent values for each individual across sessions.

Having observed conserved conduction latencies in motor and somatosensory pathways following marathon running, we next asked whether this functional stability extends to other long-range sensory systems. To address this, we measured visual and auditory evoked potentials^11,12^ before the race and within 48 hours after marathon completion. Visual evoked potentials elicited by pattern-reversal stimulation showed stable P100 peak latencies across sessions, with comparable values obtained before and after the race and no increase in inter-individual variability. Likewise, auditory evoked potentials revealed unchanged peak latencies across brainstem and cortical components, reflecting preserved signal transmission along the cochlear nerve, brainstem auditory nuclei, thalamus, and auditory cortex. Across both sensory modalities, no systematic shifts in latency were detected following endurance exercise (Fig. S1).

To further assess whether endurance exercise affects the strength of neural signal transmission, we examined the amplitudes of motor and sensory evoked potentials before and after marathon running (Fig. S2). Across all tested pathways, evoked potential amplitudes remained unchanged between the pre-run and post-run sessions, with no consistent differences observed between hemispheres.

The preservation of response amplitudes is consistent with maintained response magnitude and argues against a broad reduction in evoked responsiveness under the recording conditions. Together with the preserved conduction latencies reported above, these findings demonstrate that transient myelin remodeling occurs without measurable impairment of either the timing or the magnitude of neural signal transmission.

Although the neurophysiology cohort was necessarily small, the absence of directionally consistent latency or amplitude changes across multiple independent pathways argues against a generalized perturbation of signal timing.

### Selective and transient adaptation of higher-order cognition after marathon running

We next examined whether adaptive myelin remodeling and maintained neural signal transmission are accompanied by changes at the level of cognitive function. Cognitive performance was assessed across domains differing in their reliance on rapid information processing, interference control, and executive flexibility. This approach allowed us to determine whether endurance exercise produces global cognitive disruption or instead induces selective, transient modulation consistent with adaptive brain responses to extreme metabolic demand.

#### Processing speed (SDMT)

Processing speed was assessed using the Symbol Digit Modalities Test (SDMT)^13,14^. Mixed-effects analysis revealed a significant main effect of time on SDMT performance (F(1.98, 128.6) = 11.06, p < 0.0001), indicating changes in performance across sessions. Neither the main effect of group (F(1, 69) = 2.98, p = 0.089) nor the time × group interaction (F(2, 130) = 2.44, p = 0.091) reached statistical significance, indicating broadly similar temporal profiles across groups.

Post hoc comparisons nevertheless revealed distinct within-group patterns (Fig. 2). Non-running controls exhibited a significant improvement from baseline to the post-session assessment (p = 0.009), consistent with a robust practice effect. Marathon runners showed a smaller increase in SDMT performance immediately after the race, which increased further by the one-month follow-up, reaching statistical significance relative to baseline (p = 0.0004) and relative to the post-session (p = 0.01), approximating the magnitude of improvement observed in controls.

**Figure 2.**
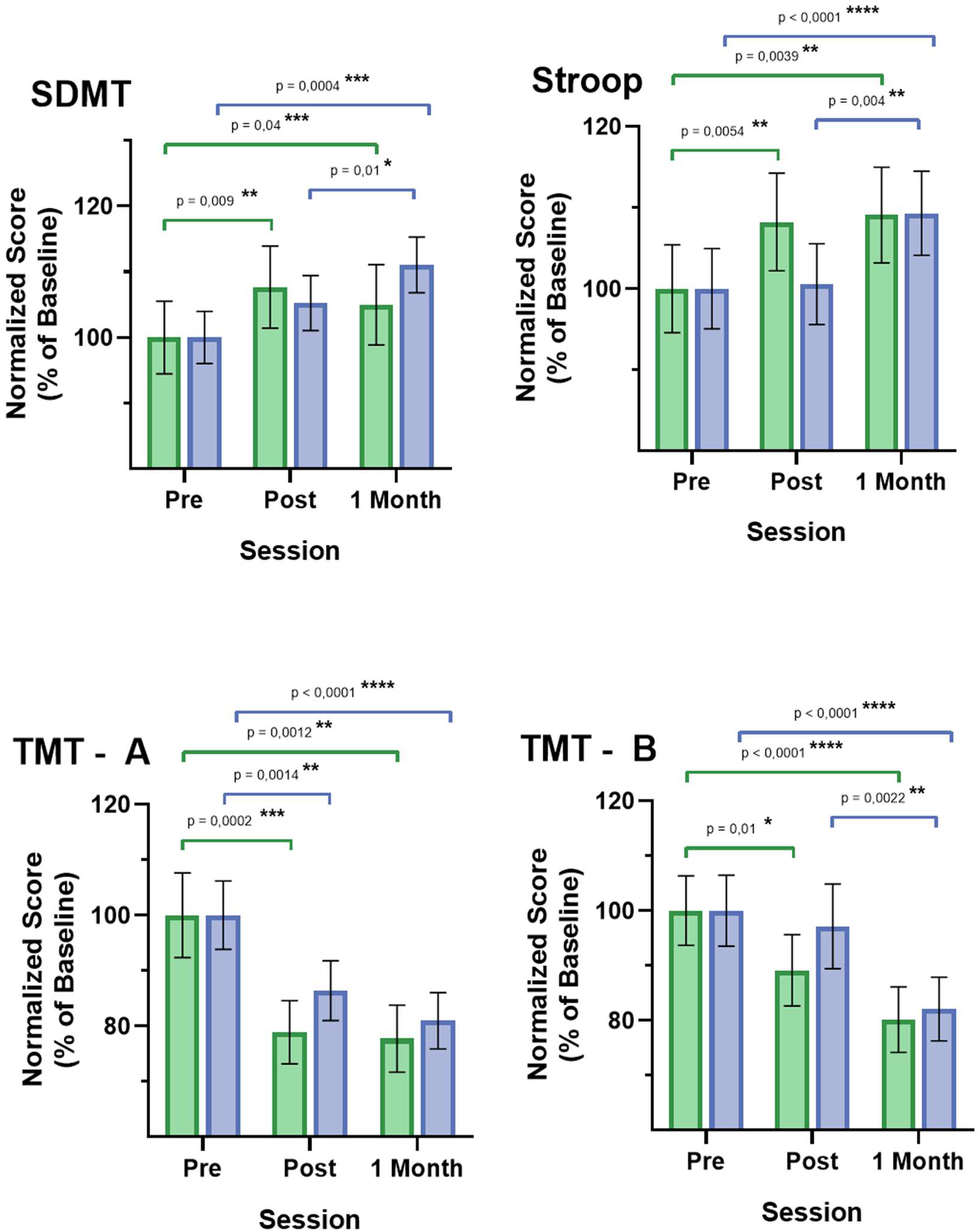
Selective and reversible modulation of cognitive performance after endurance exercise. Marathon runners (blue) and non-running controls (green) were assessed before the race (Pre), within 48 h after race completion (Post), and one month later (1 Month). Cognitive testing revealed selective and transient effects: marathon runners showed attenuated practice-related improvements in processing speed, assessed by the Symbol Digit Modalities Test (SDMT), and a transient increase in cognitive interference in the Words–Colours Stroop Test, whereas basic visuomotor speed and executive flexibility, measured by the Trail Making Test parts A and B (TMT-A, TMT-B), were preserved and exhibited robust practice effects comparable to controls. Performance across all tasks normalized by the one-month follow-up. Bars represent mean ± SEM, normalized to baseline (n = 43 for runners and n = 29 for controls). Statistical significance was assessed using Tukey’s multiple-comparisons test. Paired mean differences and 95% confidence intervals are shown in Supplementary Fig. 2.

Thus, endurance exercise did not produce a decrement in processing speed but transiently attenuated the expression of expected practice-related gains immediately after the marathon. Confidence-interval analyses supported this interpretation, showing a reduced Pre–Post improvement in runners that converged with controls by one month (Fig. S3). Together, these data indicate preserved learning capacity with short-lived modulation of processing efficiency following extreme endurance exercise.

#### Cognitive interference (Stroop)

Cognitive interference and inhibitory control were evaluated using the Words–Colours Stroop Test^15,16^. Mixed-effects analysis revealed a significant main effect of time on Stroop interference scores (F(1.70, 107.1) = 12.07, p < 0.0001), while neither the main effect of group (p = 0.33) nor the time × group interaction (p = 0.061) reached statistical significance.

Post hoc analyses showed that non-running controls exhibited a significant reduction in interference from baseline to the post-session assessment (p ≤ 0.005), consistent with practice-related improvement maintained at one month (Fig. 2). In contrast, marathon runners displayed a transient increase in interference immediately after the race, followed by normalization and subsequent improvement at the one-month follow-up. Confidence-interval analyses confirmed opposite directions of Pre–Post change in controls and runners, with convergence by one month (Fig. S3). These results indicate a short-lived, reversible modulation of interference control following endurance exercise rather than a sustained impairment of inhibitory function.

#### Visuomotor speed and executive flexibility (TMT-A and TMT-B)

Visuomotor speed and attention were assessed using the Trail Making Test, part A (TMT-A), and executive flexibility and set-shifting using part B (TMT-B)^17,18^.

For TMT-A, mixed-effects analysis revealed significant main effects of time (F(1.77, 113.6) = 31.05, p < 0.0001) and group (F(1, 69) = 8.04, p = 0.006), indicating robust improvement across sessions and overall longer completion times in marathon runners compared with controls. The time × group interaction was not significant (p = 0.61), indicating comparable temporal trajectories. Tukey-adjusted post hoc comparisons showed significant reductions in completion time from baseline to post-race and to one month in both groups (Fig. 2). Confidence-interval analyses demonstrated substantial overlap between groups across all contrasts, with no evidence of post-race slowing or delayed recovery in runners (Fig. S3).

For TMT-B, mixed-effects analysis revealed a significant main effect of time (F(2, 122) = 26.52, p < 0.0001), while neither the main effect of group (p = 0.23) nor the time × group interaction (p = 0.63) reached significance. Both groups showed practice-related improvement across sessions, although confidence-interval analyses suggested subtle differences in the temporal distribution of gains, without evidence of post-race impairment or delayed recovery in runners (Fig. S3).

Together, TMT-A and TMT-B results indicate preserved visuomotor speed and executive flexibility following marathon running. While minor variation in the timing of practice-related improvement cannot be excluded for higher-order set-shifting, these effects are modest, reversible, and do not indicate loss of executive function.

In sum, although marathon runners exhibited slightly longer completion times overall, both groups showed parallel practice-related improvements over time, reflected in overlapping confidence intervals and comparable trajectories. Because Time × Group interactions did not reach conventional significance for SDMT and Stroop, these patterns should be interpreted as evidence of selective, reversible modulation that merits confirmation in larger cohorts.

#### Synthesis

Across cognitive domains, endurance exercise was associated with a selective and transient modulation of higher-order cognitive efficiency rather than a generalized cognitive impairment. The most sensitive effects were observed in tasks placing high demands on rapid information processing and interference control, as reflected by attenuated practice-related gains in the Symbol Digit Modalities Test and a short-lived increase in Stroop interference following the marathon. These changes were reversible, with full normalization by one month. In contrast, visuomotor speed and executive flexibility, assessed using the Trail Making Test, were largely preserved: both runners and controls showed parallel practice-related improvements over time, with no evidence of delayed recovery or sustained impairment. Thus, endurance exercise preferentially affects cognitive operations requiring rapid integration and control under energetic constraint, while core visuomotor processing and set-shifting remain robust. When considered alongside conserved neural signal transmission across motor and sensory pathways, these findings support a dissociation between reversible cognitive modulation and preserved neural conduction, consistent with an adaptive physiological response to extreme metabolic demand.

## Discussion

Together, the dissociation between conserved neural signal transmission and transient cognitive adaptation is summarized in a conceptual model (Fig. 3) in which reversible myelin remodeling accompanies brain function during extreme metabolic stress. Here, the term adaptive is used to denote a reversible, non-injurious physiological response; causality and functional benefit remain to be established.

**Figure 3.**
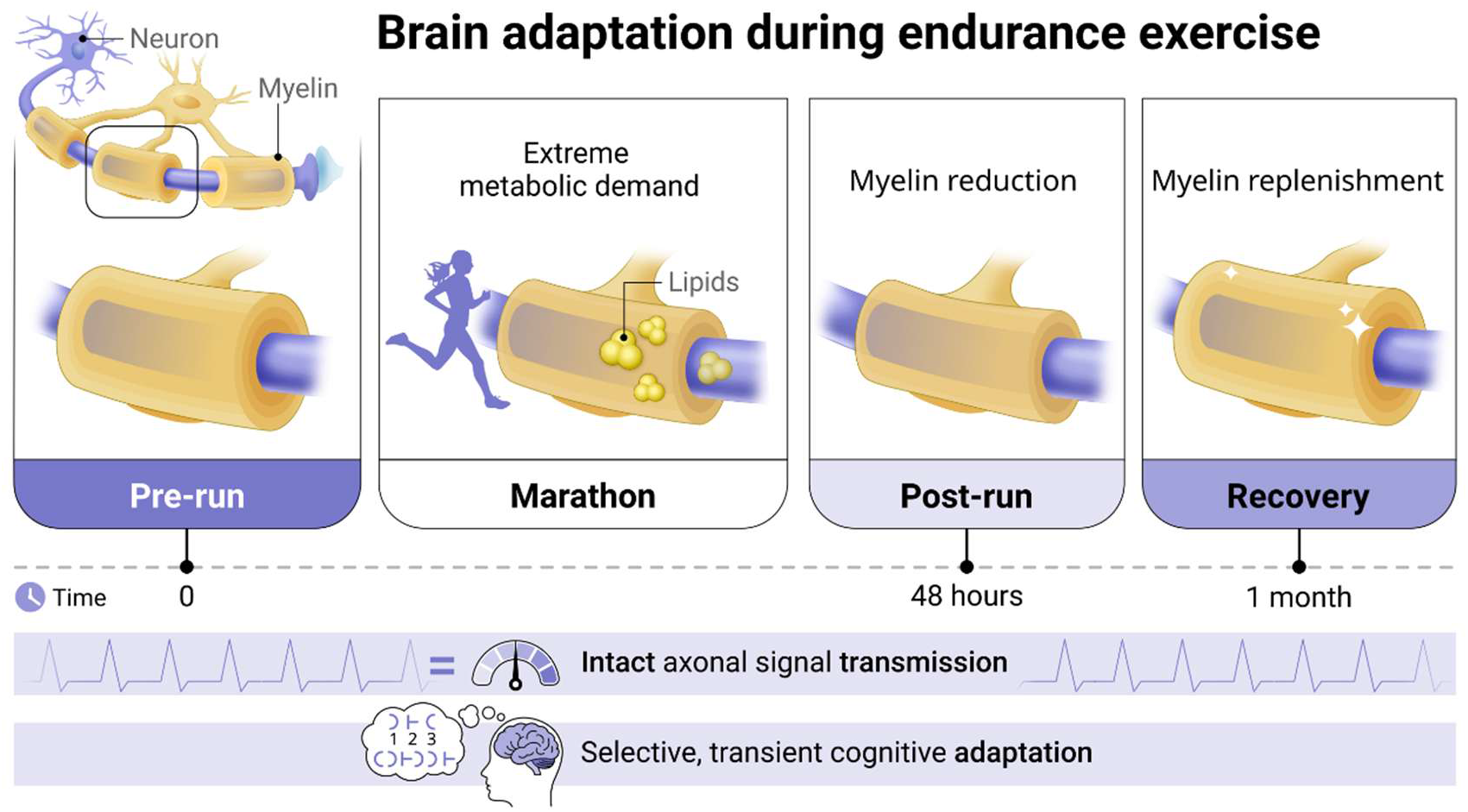
Brain adaptation during endurance exercise. Conceptual model integrating prior structural findings with the neurophysiological and cognitive results reported here. Endurance exercise imposes extreme metabolic demand on the adult human brain and is associated with reversible myelin remodeling previously demonstrated by MRI^7^. Despite transient reductions in myelin content observed immediately after marathon running, axonal signal transmission across motor and sensory pathways remains stable, as indicated by unchanged conduction latencies. During this period, cognitive effects are selective and transient, reflecting cognitive adaptation characterized by attenuated practice-related improvements in processing speed and interference control, while visuomotor speed and executive flexibility remain intact. Myelin replenishment accompanies recovery, with full normalization of cognitive performance within one month. Together, these findings indicate that reversible myelin remodeling during endurance exercise is compatible with stable neural signaling and largely maintained cognitive function, with short-lived modulation of higher-order cognitive efficiency under extreme physiological stress.

Endurance exercise provides a powerful physiological model to examine how the adult human brain responds to extreme metabolic stress. During marathon running, systemic energy demands progressively shift substrate utilization from carbohydrates toward lipids, a transition that also challenges cerebral energy homeostasis^19^. Unlike pathological conditions, this context allows interrogation of metabolic adaptations in an intact human brain operating near its physiological limits. Our findings show that this challenge is associated with a functional profile in which previously reported, reversible myelin remodeling^7^ co-occurs with conserved neural signal transmission and selective, reversible modulation of higher-order cognitive efficiency. The congruence between intact conduction across motor and sensory systems and short-lived alterations in specific cognitive domains indicates that endurance exercise engages physiological responses that maintain core neural signaling while permitting flexible adjustment of cognitive performance under energetic constraint.

A central finding of the present study is that neural signal transmission remains functionally intact despite prior evidence of transient myelin content reduction following endurance exercise^7^. Conduction latencies across multiple motor, somatosensory, visual, and auditory pathways were unchanged, indicating stable timing of long-range signal propagation. Although evoked potential latencies do not isolate conduction within specific remodeled tracts, the convergence of results across modalities argues against a broad impairment in axonal signal timing. These observations challenge the classical assumption that myelin thickness directly and linearly determines conduction velocity^8^, instead supporting the view that axonal transmission operates within a buffered functional range in mature, intact circuits, allowing short-term fluctuations in myelin content without compromising essential neural communication.

Selective modulation of higher-order cognition under extreme physiological stress is not unique to endurance exercise. Acute stress, fatigue, and hypoxic exposure have been shown to preferentially affect memory and prefrontal cortex–dependent functions while sparing motor performance, basic sensory processing, and survival-relevant behaviors^20–22^. These effects are commonly interpreted as resource-allocation strategies that prioritize essential functions during physiological challenge. Our findings extend this framework to prolonged endurance exercise, demonstrating a similar pattern of selective and reversible cognitive modulation occurring alongside intact neural signal transmission. Importantly, endurance exercise provides a physiological context in which such effects arise in the absence of overt pathology, underscoring the brain’s capacity for functional resilience under extreme metabolic demand.

Despite growing interest in this area, the cognitive consequences of endurance exercise under extreme metabolic stress remain incompletely characterized, particularly with respect to interactions between systemic metabolic signals and neural adaptation. A recent study reported reduced working-memory performance immediately after marathon completion in trained female athletes, with effects linked to markers of low energy availability^23^. Although differing in timing and cognitive demands, these observations are consistent with the notion that endurance exercise is associated with transient, energy-sensitive modulation of higher-order cognition rather than global cognitive impairment. Emerging evidence further suggests that circulating metabolic factors may contribute to such effects; for example, endurance exercise–induced signals such as lactate have been shown to influence cognitive performance and neural activity under conditions of stress and energetic challenge^24^. While circulating metabolites were not measured in the present study, our findings are compatible with this framework, demonstrating selective and reversible cognitive changes occurring alongside preserved neural signal transmission.

Consistent with this interpretation, the cognitive effects of marathon running observed here were modest, selective, and fully reversible. Changes were confined to tasks placing high demands on rapid information processing and interference control, as reflected by attenuated practice-related improvements in the SDMT and a transient increase in Stroop interference shortly after the race. These effects did not reflect absolute declines in performance but rather short-lived modulations of cognitive efficiency.

By contrast, visuomotor speed and executive flexibility, assessed with the Trail Making Test, were maintained. Both marathon runners and non-running controls exhibited robust practice effects in TMT-A and TMT-B, with parallel trajectories and overlapping confidence intervals across sessions. These findings argue against generalized cognitive slowing or fatigue-related impairment and indicate that core visuomotor processing and executive set-shifting remain intact even under extreme metabolic stress. The dissociation between higher-order cognitive efficiency and sustained visuomotor and executive function suggests that cognitive operations requiring rapid integration and interference control may be particularly sensitive to acute metabolic constraints, whereas foundational executive and sensorimotor functions remain robust.

The SDMT results further refine this interpretation. Rather than exhibiting a loss of processing speed, marathon runners showed a selective attenuation of learning-related gains immediately after the race. Confidence-interval analyses confirmed the absence of a robust post-race practice effect in runners, while performance converged fully with controls by the one-month follow-up. Such transient modulation is consistent with temporary metabolic constraints on cognitive efficiency rather than structural or functional impairment. Together with the Stroop findings, these observations indicate that endurance exercise is associated with reversible adjustments in higher-order cognitive performance that parallel intact neural conduction.

Importantly, although our data establish a close temporal association between previously reported reversible myelin remodeling, conserved conduction timing, and selective cognitive modulation, they do not demonstrate a direct causal role for myelin dynamics in sustaining neural function. Accordingly, any proposed metabolic contribution of myelin remains a working hypothesis.

Recent mechanistic work supports the plausibility of lipid-based metabolic flexibility in the nervous system under conditions of high cognitive demand. In Drosophila, memory formed after intensive training depends on neuronal mitochondrial fatty-acid β-oxidation, with cortex glia supplying lipids required to sustain this process^25^. Although Drosophila lacks myelin, these findings provide proof-of-principle that glia-derived lipids can support neuronal energy demands during cognitively intensive states and that neurons can engage fatty-acid metabolism in vivo when energetic demand is high. In this context, our observation that reversible myelin remodeling accompanies preserved neural conduction and selective cognitive modulation during endurance exercise is compatible with a broader framework in which lipid-rich glial structures, including myelin in the mammalian brain, participate in metabolic adaptation under extreme physiological demand, without implying a direct causal mechanism in humans.

Taken together, the dissociation between conserved neural signal transmission and selective cognitive modulation supports a dynamic view of myelin function in the adult brain. Rather than acting solely as a static electrical insulator, myelin appears capable of reversible remodeling that occurs alongside largely maintained brain function during periods of heightened energetic demand. Given that myelin is among the most lipid-rich biological membranes, with lipids comprising approximately 70–80% of its dry weight^5^, it represents a strategically positioned structure that may participate in metabolic adaptation, analogous to peripheral fat stores mobilized during prolonged exercise^19^. Within this framework, transient myelin remodeling may accompany redistribution of energetic resources while neural signaling fidelity is maintained.

These findings have broader implications for brain resilience across the lifespan and in disease. The capacity to dynamically regulate myelin under metabolic stress may contribute to neural robustness in healthy adults, whereas impairment of such processes could increase vulnerability in aging or neurological disorders characterized by disrupted energy metabolism or myelin integrity^26^.

Several limitations should be acknowledged. The intensive nature of repeated neurophysiological and cognitive testing in marathon runners constrained sample size, positioning this work as a proof-of-principle study in human physiology. Nonetheless, the internal consistency across independent neurophysiological and cognitive measures strengthens the robustness of the conclusions. Future studies combining metabolic profiling, electrophysiology, and longitudinal cognitive assessment will be required to directly link energy availability, myelin dynamics, and functional outcomes. Extending this approach to older individuals and patient populations will further clarify whether these adaptive responses are preserved or disrupted in disease. Finally, because neurophysiological and cognitive measures were not acquired simultaneously within individuals, future work will be needed to determine whether inter-individual variability in conduction stability relates to the magnitude of cognitive modulation.

In summary, the adult human brain tolerates extreme metabolic stress through physiological responses that conserve core neural signaling while allowing selective, transient modulation of higher-order cognitive efficiency. By integrating neurophysiological and cognitive measures with prior structural findings, our results support a dynamic view in which reversible myelin remodeling occurs within an adaptive physiological framework compatible with sustained brain function under metabolic challenge. Rather than demonstrating cognitive failure during extreme endurance exercise, these findings reveal a dissociation between intact neural signal transmission and selective, reversible modulation of higher-order cognition, highlighting an adaptive framework for brain resilience under metabolic stress.

## Acknowledgments

We thank all participants for their time, commitment, and generosity in contributing to this study. In particular, we acknowledge the marathon runners for their exceptional effort in completing demanding physiological and cognitive assessments before and after the race, and the control volunteers for their equally essential participation across repeated testing sessions. We are also grateful to the technical and clinical staff involved in data acquisition and neurophysiological recordings, and to Zigor Aira for reading the manuscript and suggestions.

## Funding

This work was supported by (all awarded to CM)

National Network of Biomedical Research in Neurodegenerative Diseases (CIBERNED, grant no. CB06/05/00 76),

The Ministerio de Ciencia, Innovación y Universidades (MCINN, grant no. PID2022-143020OB-I00),

The Gobierno Vasco (grant no. IT1551-22), and EITB-Maratoia (grants no. BIO22/ALZ/015 and BIO23/EM/002).

E.L.-P. holds a fellowship from Gobierno Vasco.

The funders had no role in study design, data collection and analysis, decision to publish, or preparation of the manuscript.

## Author contributions

Conceptualization: CM

Methodology: IL, SP, IYS, ELM, GGG, CM

Investigation: IL, SP, IYS, ELM, GGG, CM

Visualization: IL, SP, IYS, ELM, GGG, CM

Funding acquisition: CM

Project administration: CM

Supervision: CM

Writing – original draft: CM

Writing – review & editing: IL, SP, IYS, ELM, GGG, CM

## Competing interests

Authors declare that they have no competing interests.

## Data and materials availability

All data are available in the main text or the supplementary materials.

## Supplementary Materials

## Materials and Methods

### Participant recruitment

Healthy adult volunteers were recruited and assigned to either a running group or a non-running control group. Marathon runners were recruited among registered participants of officially organized marathon or ultramarathon, and had completed systematic endurance training in preparation for the race. Inclusion criteria for runners included the absence of neurological, psychiatric, cardiovascular, or metabolic disorders, and the ability to complete the marathon without medical complications. Control participants were recruited from the same population and were matched to runners for age and sex as closely as possible. Controls did not participate in endurance events during the study period and maintained their usual daily activities.

All participants were assessed at three time points: before the race (Pre), within 48 hours after marathon completion (Post), and one month later (1 Month). Control participants were tested at comparable intervals to account for practice effects and time-dependent variability. Participants were instructed to refrain from strenuous physical activity, alcohol consumption, and caffeine intake for at least 24 hours before each testing session.

Exclusion criteria for all participants included a history of neurological disease, head injury, substance abuse, use of medications affecting the central nervous system, or contraindications to neurophysiological testing. All participants provided written consent prior to participation. The study was conducted in accordance with the Declaration of Helsinki and was approved by the Ethics Committee for Research of Euskadi.

### Evoked potential recordings, electrode montages, and signal processing

Evoked potentials were recorded to assess the integrity and timing of neural signal transmission across motor and sensory pathways before and after marathon running. All recordings were performed using standard clinical neurophysiology procedures under identical experimental conditions across sessions, enabling within-subject comparisons of conduction latency over time. Scalp electrodes were positioned according to the international 10–20 system, impedances were kept below 5 kΩ, and signals were amplified and digitized using clinical-grade acquisition systems.

Motor evoked potentials (MEPs) elicited by transcranial magnetic stimulation (TMS) of the primary motor cortex were used as a functional readout of corticospinal signal transmission, with response latency reflecting conduction timing to upper- and lower-limb muscles^9^. Single-pulse TMS was delivered over M1 using a figure-of-eight coil positioned tangentially to the scalp, inducing a focal intracortical current that generated descending action potentials along the corticospinal tract. MEPs were recorded from target muscles innervated by the median and tibial nerves using bipolar surface electromyography electrodes placed over the muscle belly with a tendon reference. EMG signals were band-pass filtered at 10–1,000 Hz, sampled at ≥5 kHz, and latency was defined as the interval between the TMS trigger and EMG onset.

Somatosensory evoked potentials elicited by peripheral nerve stimulation were used as a functional readout of ascending sensory conduction, with peak latencies reflecting the timing and integrity of dorsal column–medial lemniscal pathway transmission to primary somatosensory cortex^10^. SEPs were elicited by electrical stimulation of the median nerve at the wrist and the posterior tibial nerve at the ankle. Cortical responses were recorded using CP3–Fz (right median nerve), CP4 –Fz (left median nerve) and CPz–Fz (posterior tibial nerve) montages. Signals were band-pass filtered at 5– 2,000 Hz, sampled at ≥5 kHz, averaged across repeated stimuli, and peak latencies of the N20 (upper limb) and P37 (lower limb) components were measured as indices of conduction timing along ascending somatosensory pathways.

Pattern-reversal visual evoked potentials provide a robust measure of visual pathway conduction, with the P100 peak latency reflecting the timing of signal transmission from the retina to primary visual cortex^11^. VEPs were recorded in response to pattern-reversal checkerboard stimulation. Cortical responses were obtained from Oz referenced to Fz (Oz–Fz). Signals were band-pass filtered at 1–100 Hz, sampled at ≥1 kHz, averaged across trials, and the P100 peak latency was quantified.

Auditory evoked potentials were used as a functional readout of auditory pathway integrity and conduction timing, with component latencies reflecting signal transmission from the cochlear nerve through brainstem relays and, for longer-latency responses, thalamocortical and cortical processing^12^. In particular, auditory brainstem responses provide a robust measure of auditory pathway conduction, with the latency of Wave V reflecting signal transmission through the rostral brainstem and serving as a reliable index of auditory brainstem integrity. AEPs were elicited using binaural auditory stimulation, and responses were recorded using a Cz–A1/A2 montage. Brainstem auditory responses were band-pass filtered at 100–3,000 Hz and sampled at ≥10 kHz, whereas middle-latency and cortical auditory responses were filtered at 1–100 Hz and sampled at ≥1 kHz. Peak latencies of canonical auditory components were measured to assess conduction timing along the auditory pathway.

Across all modalities, stimulation parameters, electrode placement, and acquisition settings were kept constant within each participant across sessions. Latency measures were selected a priori as the primary outcome because they provide a robust and modality-independent index of axonal signal transmission that is minimally influenced by short-term fluctuations in excitability or attention.

### Neuropsychological assessment

Neuropsychological testing was performed to evaluate cognitive domains potentially sensitive to endurance exercise and metabolic stress, with a focus on processing speed, cognitive interference, visuomotor performance, and executive flexibility. Standardized tests commonly used in clinical and research settings were administered to marathon runners and non-running controls at three time points: before the race (Pre), within 48 hours after race completion (Post), and one month later (1 Month). All tests were administered in a standardized manner by trained personnel. A fixed test order was used across sessions to minimize procedural variability among subjects.

Processing speed was assessed using the Symbol Digit Modalities Test (SDMT), a widely used timed substitution task in which participants match symbols to digits according to a key; total correct responses within the time limit provide a validated index of cognitive processing speed^13,14^. Participants were required to match symbols to numbers according to a reference key as quickly and accurately as possible within a fixed time period. The primary outcome measure was the total number of correct responses, providing an index of rapid information processing and attentional efficiency.

Cognitive interference and control were evaluated using the Words–Colours Stroop Test, in which participants name the ink color of colored words printed in congruent or incongruent hues. Performance in the incongruent condition requires suppression of the automatic word-reading response, and the resulting interference score reflects the ability to inhibit cognitive interference and exert executive control^15,16^. Participants completed conditions requiring reading of words, naming of colors, and naming of ink colors incongruent with the written word. The interference score, derived from performance in the incongruent condition relative to baseline conditions, was used as a measure of susceptibility to cognitive interference and executive control.

Visuomotor speed and attention were assessed with the Trail Making Test, part A (TMT-A), in which participants connect numbered circles in ascending order as quickly and accurately as possible. Completion time in TMT-A provides a validated index of visuomotor scanning, processing speed, and sustained attention^17^. Participants were instructed to connect numbered circles in ascending order as quickly as possible. Completion time was recorded as the primary outcome measure.

Executive flexibility and set-shifting were assessed using the Trail Making Test, part B (TMT-B), in which participants alternate between numbers and letters in ascending order as quickly and accurately as possible. Completion time in TMT-B provides a validated index of cognitive flexibility and executive control beyond basic processing speed^18^. In this condition, participants alternated between numbers and letters in ascending order. Completion time was recorded as an index of executive function and cognitive flexibility.

For all tests, raw scores or completion times were used for statistical analyses as specified in the Results. Tests were selected to allow differentiation between higher-order cognitive processes sensitive to metabolic constraints and more basic visuomotor or executive functions. Repeated testing effects were accounted for by comparing performance trajectories between runners and controls across sessions.

### Statistical analysis

Statistical analyses were performed using Graph Pad statistical software. Data are presented as mean ± SEM unless otherwise indicated. For neurophysiological measures, latency values were analyzed using within-subject comparisons across time points (Pre, Post, and, where available, 1 Month). Given the proof-of-principle nature of the neurophysiology experiments and the limited sample size, analyses focused on descriptive statistics and consistency of effects across pathways rather than formal hypothesis testing.

For neuropsychological measures, group differences across time were assessed using linear mixed-effects models (REML) with time and group as fixed effects and subject as a random effect to account for repeated measurements. Practice effects were evaluated via the main effect of time, and potential group differences in performance trajectories were assessed using the Time × Group interaction. When omnibus effects were significant, Tukey-adjusted post-hoc comparisons were conducted to explore within-group changes across sessions. To satisfy model assumptions, log_10_ transformation was applied to TMT-B completion times; all other measures were analysed on their original scale. A significance threshold of *P* < 0.05 was applied.

Pretreatment of the data was performed to eliminate outliers using the interquartile (IQR) method, discarding all data below Q1 – 1.5 × IQR or over Q3 + 1.5 × IQR limits^27,28^. Model assumptions of normality of residuals, homocedasticity and linearity of fixed effects were assessed qualitatively based on data distribution and consistency across measures. Given the exploratory and integrative nature of the study, emphasis was placed on the convergence of results across independent cognitive domains and neurophysiological pathways. Exact *P* values are reported where relevant.

---

**Supplementary Figure 1.**
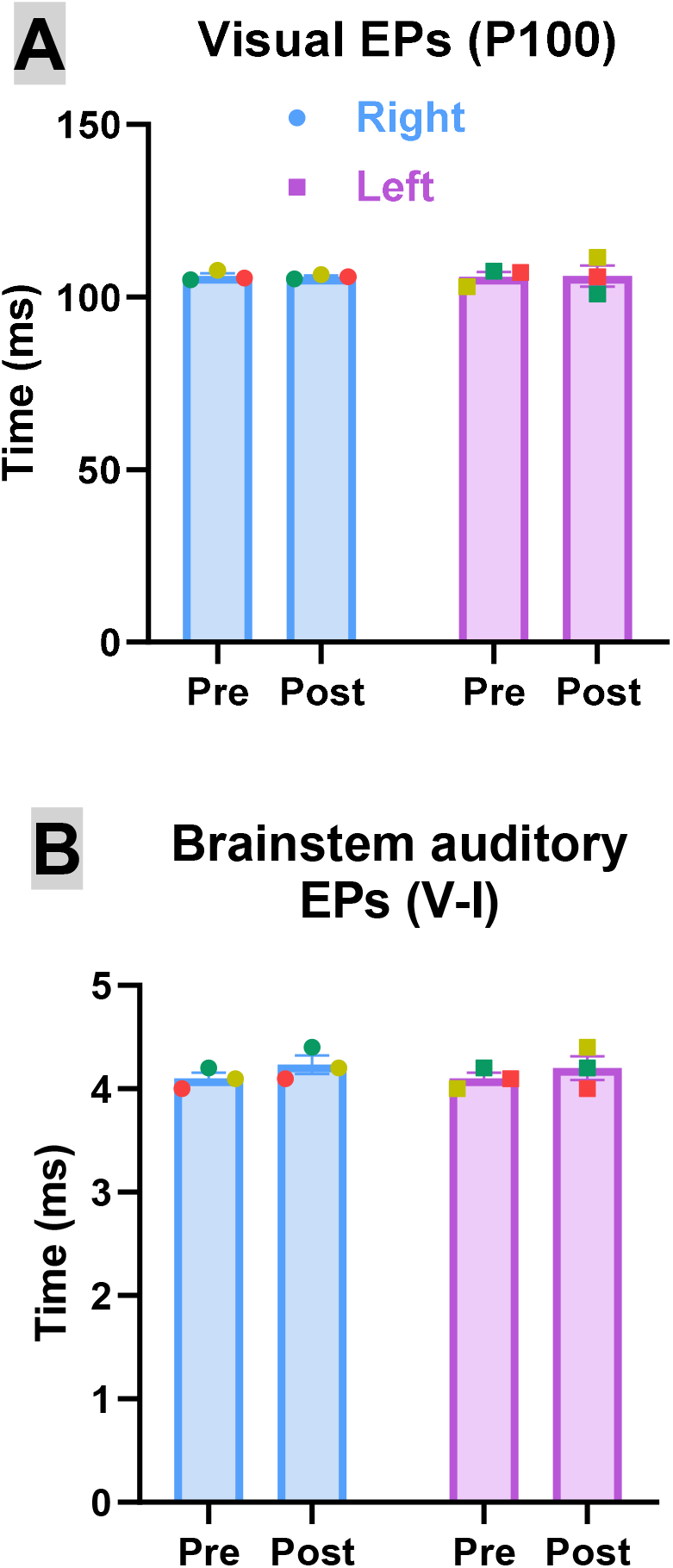
Visual and auditory evoked potential latencies are intact after endurance exercise. Representative visual and auditory evoked potentials (EPs) recorded before the race (Pre) and within 48 hours after marathon completion (Post). (A) Visual EPs show the canonical P100 component, reflecting conduction along the visual pathway, (B) while brainstem auditory EPs (V-I) display components reflecting conduction along the auditory pathway. Quantification of peak latencies revealed no differences between sessions for either modality and no increase in variability, indicating unaffected conduction timing in visual and auditory systems following endurance exercise. Values are mean ± SEM (n=3), no detectable changes, paired two-tailed t test, in all comparisons. EPs were recorded bilaterally, in the right and left hemispheres.

**Supplementary Figure 2.**
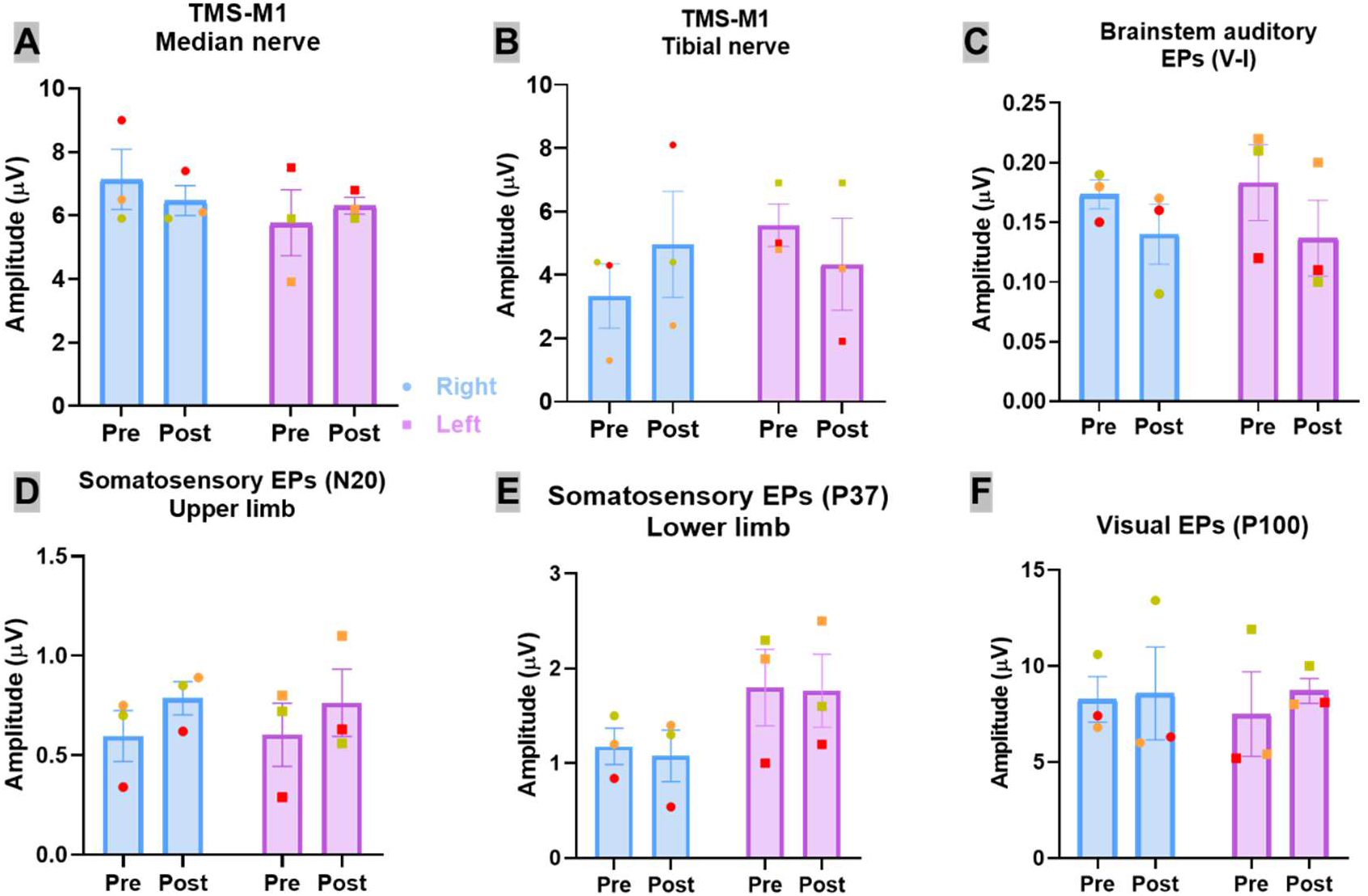
Evoked potential amplitudes are stable after endurance exercise. Peak-to-peak amplitudes of motor evoked potentials (MEPs; A–B) and sensory evoked potentials (SEPs; C–F) recorded before the marathon (Pre) and within 48 h after race completion (Post). Responses are shown separately for right (blue) and left (purple) stimulation/recordings, with individual data points overlaid on group means ± SEM. Across all modalities and hemispheres, evoked potential amplitudes remained stable between sessions, with no systematic reductions or increases following endurance exercise. These data indicate unaffected synaptic and axonal excitability and complement latency measures, supporting intact neural signal transmission despite transient myelin remodeling. N=3, P > 0.5, paired two-tailed t test, in all comparisons.

**Supplementary Figure 3.**
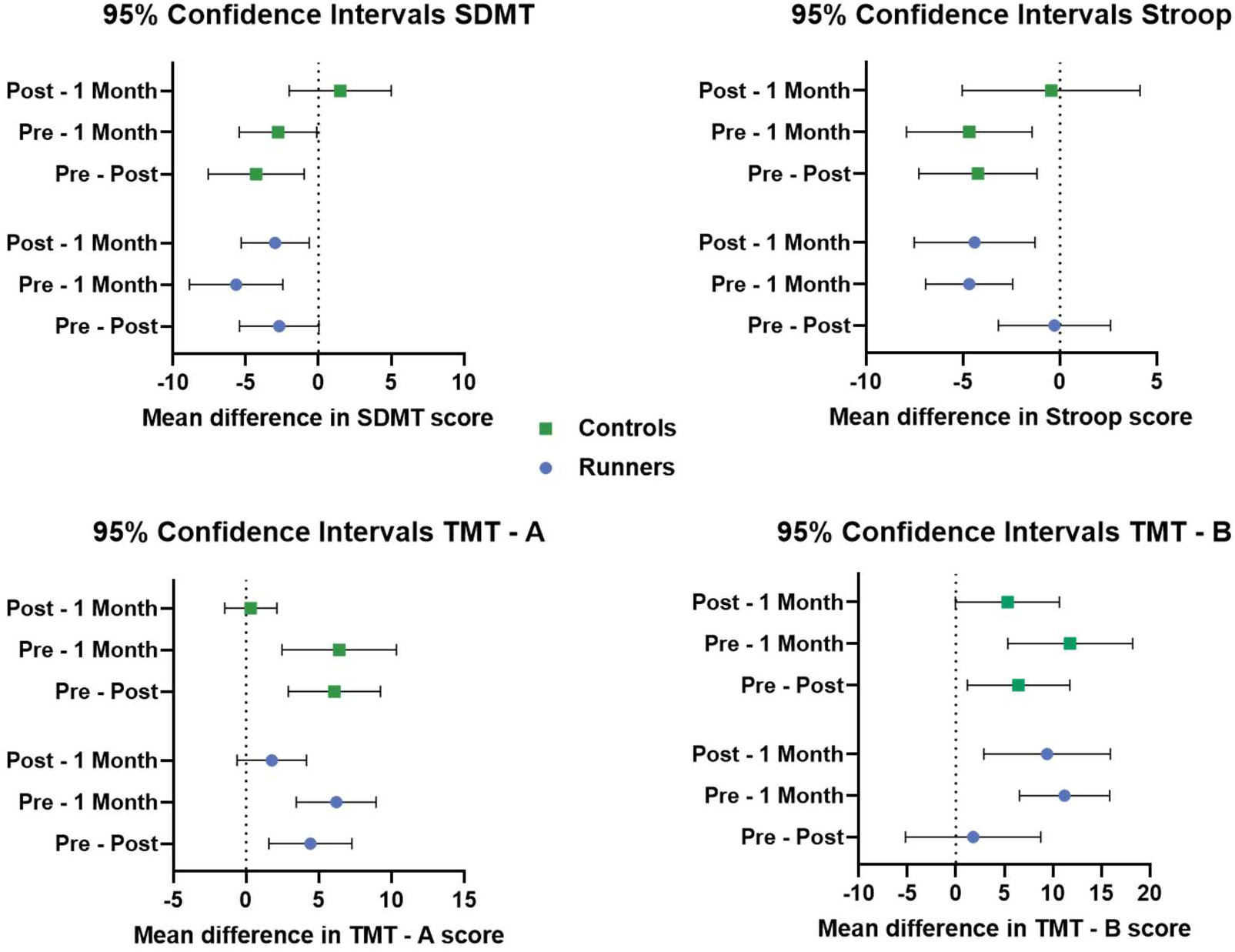
Paired mean differences and confidence intervals for cognitive performance across sessions. Forest plots showing paired mean differences (±95% confidence intervals) for changes in cognitive performance across sessions in marathon runners (blue) and non-running controls (green). Comparisons include Pre–Post (within 48 h after the race), Pre–1 Month, and Post–1 Month intervals for the Symbol Digit Modalities Test (SDMT), Words–Colours Stroop Test, Trail Making Test part A (TMT-A), and Trail Making Test part B (TMT-B). The vertical dashed line denotes no change. For SDMT and Stroop, runners show attenuated or transiently reversed practice-related gains immediately post-race relative to controls, with confidence intervals converging by one month. In contrast, for TMT-A and TMT-B, both groups exhibit comparable improvements across sessions, with overlapping confidence intervals and no evidence of delayed recovery or increased variability in runners. These interval estimates complement the group-level analyses shown in Fig. 2 and highlight the selectivity and reversibility of cognitive modulation following endurance exercise.

## Notes

### Competing Interest Statement

The authors have declared no competing interest.

## References and Notes

1. Hawley, J. A., Lundby, C., Cotter, J. D. & Burke, L. M. Maximizing Cellular Adaptation to Endurance Exercise in Skeletal Muscle. Cell Metab. 27, 962–976 (2018).

2. Romijn, J. A. et al. Regulation of endogenous fat and carbohydrate metabolism in relation to exercise intensity and duration. Am. J. Physiol. 265, E380–391 (1993).

3. Bolaños, J. P. & Magistretti, P. J. The neuron-astrocyte metabolic unit as a cornerstone of brain energy metabolism in health and disease. Nat. Metab. 7, 2414–2423 (2025).

4. Rojas, R., Pellitero, A., Arslan, B., Pérez-Samartín. A. & Matute, C. The Neurobiology of Fatty Acids. Prog. Lipid Res. in press, (2026).

5. Norton, W. T. & Poduslo, S. E. Myelination in rat brain: changes in myelin composition during brain maturation. J. Neurochem. 21, 759–773 (1973).

6. Asadollahi, E. et al. Oligodendroglial fatty acid metabolism as a central nervous system energy reserve. Nat. Neurosci. 27, 1934–1944 (2024).

7. Ramos-Cabrer, P. et al. Reversible reduction in brain myelin content upon marathon running. Nat. Metab. 7, 697–703 (2025).

8. Fields, R. D. A new mechanism of nervous system plasticity: activity-dependent myelination. Nat. Rev. Neurosci. 16, 756–767 (2015).

9. Spampinato, D. A., Ibanez, J., Rocchi, L. & Rothwell, J. Motor potentials evoked by transcranial magnetic stimulation: interpreting a simple measure of a complex system. J. Physiol. 601, 2827–2851 (2023).

10. Fustes, O. J. H. et al. Somatosensory evoked potentials in clinical practice: a review. Arq. Neuropsiquiatr. 79, 824–831 (2021).

11. Šuštar Habjan, M. et al. ISCEV standard for clinical visual evoked potentials (2025 update). Doc. Ophthalmol. Adv. Ophthalmol. 151, 97–112 (2025).

12. Kwak, C. et al. Understanding Standard Procedure in Auditory Brainstem Response: Importance of Normative Data. J. Audiol. Otol. 28, 243–251 (2024).

13. Smith, A. Symbol Digit Modalities Test. 10.1037/t27513-000 (2016).

14. Seo, D. et al. Digital symbol-digit modalities test with modified flexible protocols in patients with CNS demyelinating diseases. Sci. Rep. 14, 14649 (2024).

15. Stroop, J. R. Studies of interference in serial verbal reactions. J. Exp. Psychol. 18, 643–662 (1935).

16. Scarpina, F. & Tagini, S. The Stroop Color and Word Test. Front. Psychol. 8, 557 (2017).

17. Reitan, R. M. & Wolfson, D. The Trail Making Test as an initial screening procedure for neuropsychological impairment in older children. Arch. Clin. Neuropsychol. Off. J. Natl. Acad. Neuropsychol. 19, 281–288 (2004).

18. Pellas, J. & Damberg, M. Assessment of executive functions in older adults: Translation and initial validation of the Swedish version of the Frontal Assessment Battery, FAB-Swe. Appl. Neuropsychol. Adult 31, 64–68 (2024).

19. Alghannam, A. F., Ghaith, M. M. & Alhussain, M. H. Regulation of Energy Substrate Metabolism in Endurance Exercise. Int. J. Environ. Res. Public. Health 18, 4963 (2021).

20. Arnsten, A. F. T. Toward a new understanding of attention-deficit hyperactivity disorder pathophysiology: an important role for prefrontal cortex dysfunction. CNS Drugs 23 Suppl 1, 33–41 (2009).

21. Diamond, A. Executive functions. Annu. Rev. Psychol. 64, 135–168 (2013).

22. Feddersen, B. et al. Regional differences in the cerebral blood flow velocity response to hypobaric hypoxia at high altitudes. J. Cereb. Blood Flow Metab. Off. J. Int. Soc. Cereb. Blood Flow Metab. 35, 1846–1851 (2015).

23. Boere, K. et al. Working Memory Performance is Reduced Following a Marathon Race and Associated with Low Energy Availability in Females. Med. Sci. Sports Exerc. https://doi.org/10.1249/MSS.0000000000003937 (2026) doi:10.1249/MSS.0000000000003937.

24. Clairis, N., Barakat, A., Brochard, J., Xin, L. & Sandi, C. A neurometabolic mechanism involving dmPFC/dACC lactate in physical effort-based decision-making. Mol. Psychiatry 30, 899–913 (2025).

25. Pavlowsky, A. et al. Neuronal fatty acid oxidation fuels memory after intensive learning in Drosophila. Nat. Metab. 7, 2438–2450 (2025).

26. Matute, C. & Verkhratsky, A. Brain myelin as a deficient energy source in aging and disease. Trends Endocrinol. Metab. TEM 36, 781–784 (2025).

27. Grubbs, F. E. Sample Criteria for Testing Outlying Observations. Ann. Math. Stat. 21, 27–58 (1950).

28. Smiti, A. A critical overview of outlier detection methods. Comput. Sci. Rev. 38, 100306 (2020).

